# Artificial nanovesicles for dsRNA delivery in spray induced gene silencing for crop protection

**DOI:** 10.1101/2023.01.03.522662

**Authors:** Lulu Qiao, Jonatan Niño-Sánchez, Rachael Hamby, Luca Capriotti, Angela Chen, Bruno Mezzetti, Hailing Jin

## Abstract

Spray-Induced Gene Silencing (SIGS) is an innovative and eco-friendly technology where topical application of pathogen gene-targeting RNAs to plant material can enable disease control. SIGS applications remain limited because of the instability of dsRNA, which can be rapidly degraded when exposed to various environmental conditions. Inspired by the natural mechanism of crosskingdom RNAi through extracellular vesicle trafficking, we describe herein the use of artificial nanovesicles (AVs) for dsRNA encapsulation and control against the fungal pathogen, *Botrytis cinerea*. AVs were synthesized using three different cationic lipid formulations, DOTAP + PEG, DOTAP, and DODMA, and examined for their ability to protect and deliver dsRNA. All three formulations enabled dsRNA delivery and uptake by *B. cinerea*. Further, encapsulating dsRNA in AVs provided strong protection from nuclease degradation and from removal by leaf washing. This improved stability led to prolonged RNAi-mediated protection against *B. cinerea* both on pre- and post-harvest plant material using AVs. Specifically, the AVs extended the protection duration conferred by dsRNA to 10 days on tomato and grape fruits and to 21 days on grape leaves. The results of this work demonstrate how AVs can be used as a new nanocarrier to overcome dsRNA instability in SIGS for crop protection.

## Introduction

Fungal plant diseases are a major threat to global food security^1,2^, causing perennial crop yield losses of up to 20%, and postharvest product losses of up to 10% worldwide^3^. For example, *Botrytis cinerea*, the causal agent of gray mold disease in over 1000 plant species, alone causes billions of dollars in annual crop yield losses^4,5^. Alarmingly, this threat is projected to increase as rising temperatures associated with global warming favor fungal pathogen growth^2,6^. Currently, the most widely used plant pathogen control practices require routine application of fungicides which has contributed to the rapid development of fungicide resistant pathogens^3^. To safeguard global food security, an alternative, environmentally friendly fungal control method must be developed. Recent studies have shown that many aggressive fungal pathogens can take up RNAs from the environment^7–11^. These RNAs can then induce silencing of fungal genes with complementary sequences. This discovery led to the development of Spray-Induced Gene Silencing (SIGS), where fungal virulence gene-targeting double-stranded RNAs (dsRNAs) or small RNAs (sRNAs) are topically applied to plant material to control fungal pathogens.

SIGS can provide safe and powerful plant protection on both pre-harvest crops and postharvest products against fungal pathogens that have high RNA uptake efficiency^12–14^. SIGS RNAs can be versatilely designed to target non-conserved regions of the genes and to be species-specific, minimizing the risk of off-target effects on other organisms. They can also be designed to target multiple genes from the same or different pathogens simultaneously. *B. cinerea* was found to deliver sRNAs into plant cells that silenced host immunity genes by commandeering *A. thaliana* Argonaute 1 (AGO1) protein to evoke cross-kingdom RNAi^15^. These transferred *B. cinerea* sRNAs are generated by fungal Dicer-Like (DCL) proteins (*B. cinerea* DCL1 and DCL2). Studies in other organismal interactions have demonstrated other microbes/pathogens, including plant fungal pathogens *Fusarium oxysporum^18^* and *Verticillium dahliae^7^*, an oomycete pathogen *H. arabidopsidis*^16^, an animal fungal pathogen *Beauveria bassiana* of mosqiuito^17^, a bacterial pathogen^19^ and even symbiotic microbes, such as ectomycorrhizal fungus *Pisolithus microcarpus*^20^ and Rhizobium (*Bradyrhizobium japonicum*)^21^, also utilize host Argonaute proteins for cross-kingdom RNAi. The biogenesis of these pathogen sRNAs involved in cross-kingdom RNAi are also largely dependent on pathogen DCL proteins^7–8,10,15,22–26^. These studies demonstrate crosskingdom RNAi is a conserved mechanism across different species, including plant and animal hosts. Consequently, SIGS RNAs which target genes involved in RNAi machinery and in sRNA biogenesis have proven to be quite effective^7–8,10,23,25–28^, making them ideal biological pathways to target in other organisms. Furthermore, because RNAi can tolerate multiple mismatches between sRNAs and target RNAs^29^, fungal pathogens are less likely to develop resistance to SIGS RNAs than to traditional fungicides, which mostly bind to and inhibit a specific enzyme or protein. To date, SIGS has effectively been used to control a wide range of insect pests^30–33^, viruses^34–35^ and pathogenic fungi including *Fusarium graminearum* infection in barley^9,10^, and gray mold disease on fruits, vegetables and flowers^7,11,13–14,36^.

One major drawback of SIGS is the relative instability of RNA in the environment, particularly when subjected to rainfall, high humidity, or UV light^33^. Thus, improving environmental RNA stability is critical for successful SIGS applications. One strategy is to dock RNAs in synthetic inorganic materials. Specifically, dsRNAs targeting plant viruses have been loaded into layered double hydroxide (LDH) nanosheets to protect dsRNA from nuclease degradation^34–35^ and increase the stability and the durability of the RNAi effect. Ultimately, this can provide RNAi-based systemic protection against several plant viruses for at least 20 days after topical application^34,37^. This nanotechnology was recently proven to be effective for plant protection against fungal pathogens^27^.

In nature, plants and animals encapsulate RNAs in extracellular vesicles (EVs) for safe transportation between cells or interacting organisms^38–42^. Recent advances in RNA-based pharmaceuticals have utilized liposomes, synthetic spherical lipid-based nanoparticles for drug delivery, notably in both mRNA COVID-19 vaccines^43–48^. Liposome formulations are often cationic to facilitate dsRNA binding through electrostatic interactions^45,49–50^, which protects the dsRNAs from degradation. Drawing inspiration from both naturally occurring vesicles and the clinical use of liposomes, here, we package fungal gene-targeting RNAs in artificial nanovesicles (AVs), to mimic the plant’s natural RNA delivery strategy for use in SIGS applications.

In this study, we demonstrate that dsRNAs packaged in AVs can be successfully utilized in crop protection strategies. Three types of AVs were synthesized and found to confer protection to loaded dsRNA, which remained detectable in large amounts on plant surfaces over a long period of time. When applied to plants, AV-dsRNA can extend the length of fungal control conferred by fungal gene-targeting dsRNA to crops. Overall, this work demonstrates how organic nanoparticles can be utilized to strengthen SIGS-based crop protection strategies.

## Results

### Artificial nanovesicles protect and efficiently deliver dsRNAs to the fungal pathogen

#### Botrytis cinerea

PEGylated AVs were synthesized using the lipid film hydration method for cationic liposomes^49^. Specifically, AVs were generated using a mixture of the cationic lipid 1,2-dioleoyl 3-trimethylammonium-propane (DOTAP), cholesterol and 1,2-distearoyl-sn-glycero-3-phosphoethanolamine-N-[methoxy(polyethyleneglycol)-2000] (DSPE-PEG2000)^49^. This AV composition had used to deliver dsRNA to mammalian cells, so we hypothesized that it could also be ideal for antifungal RNAi applications^49^. We then established the loading ratio necessary for the AVs to completely encapsulate dsRNAs of interest. Previous studies have shown that fungal Dicerlike proteins play a critical role in the virulence of several fungal pathogens, such as *Botrytis cinerea*, *Sclerotinia sclerotiorum*, *Verticillium dahliae*, *Fusarium graminearum*, *Penicillium italicum*, and *Valsa mali*, and making them ideal targets for SIGS^7–8,10,15,22–26^. Dicer-like proteins generate fungal sRNA effectors that silence host immunity genes through cross-kingdom RNAi^7,15^. Furthermore, since pathogens have been shown to hijack host Argonaute proteins using sRNA effectors for cross-kingdom RNAi in multiple systems, Dicer-like proteins are strong candidates for inhibiting pathogenicity and virulence. Exogenous treatment of *Bc-DCL1/2-dsRNA*, a dsRNA integrating fragments of the Dicer-like 1 (252 bp) and Dicer-like 2 (238bp) sequences from *Botrytis cinerea*, on the plant leaf surface can efficiently inhibit fungal disease^7^. When loading AVs, the loading ratio is determined by the N:P ratio, where N = # of positively charged polymer nitrogen groups and P = # of negatively charged nucleic acid phosphate groups. Here, DOTAP has one positively charged nitrogen per molecule, whereas the double stranded *Bc-DCL1/2* RNA, which is 490 nucleotides in length, has 980 negatively charged nucleic acid groups per molecule. Thus, loading ratios between the AVs and the *Bc-DCL1/2-dsRNA*, from 1:1 to 4:1, were examined to identify the minimum amount of AVs required to bind all the dsRNA present in the solution. We concluded that a 4:1 (AV:dsRNA) ratio was the optimal ratio needed for dsRNA loading as *Bc-DCL1/2-dsRNA* loaded into AVs at this ratio could not migrate from the loading well due to complete association with the AVs (Figure 1A). The average size, polydispersity index (PDI), and zeta potential (ZP) of the loaded AVs was also determined. The AV-*Bc-DCL1/2*-dsRNA lipoplexes had an average size of 365.3 nm with PDI = 0.455 and ZP = +47.17 mV (Table S1).

**Figure 1.**
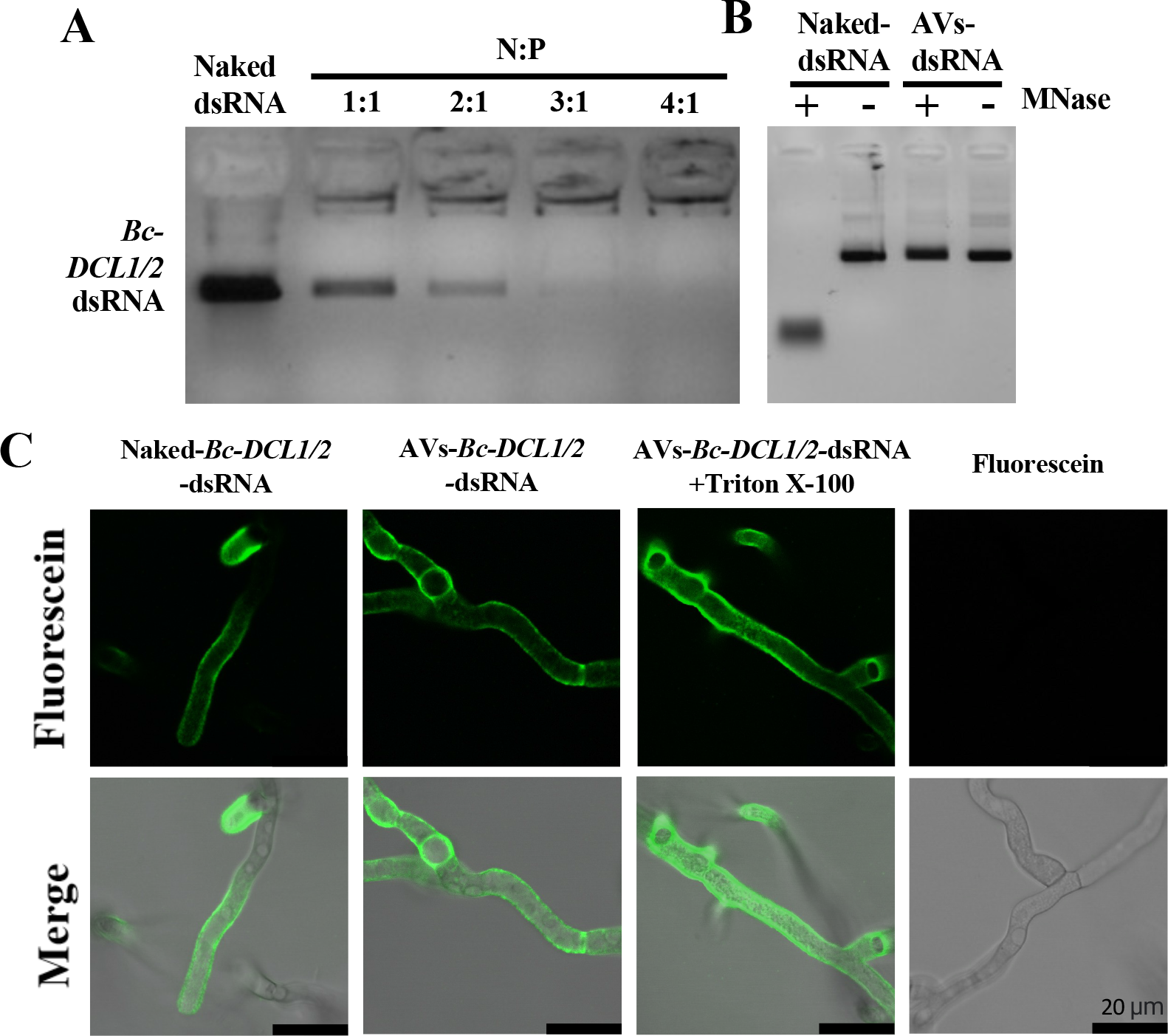
dsRNA loaded into AVs is shielded from nuclease degradation and easily taken up by *Botrytis cinerea*. (**A**) AV-*Bc-DCL1/2*-dsRNA lipoplexes were formed at a range of indicated charge ratios (N:P) and incubated for 2 h at room temperature before being loaded onto a 2% agarose gel. Complete loading was achieved at an AV:dsRNA mass ratio of 4:1. (**B**) The stability of naked- and AV-*Bc-DCL1/2*-dsRNA was tested after MNase treatment. *Bc-DCL1/2*-dsRNA was released from AVs using 1% Triton X-100 before gel electrophoresis. (**C**) Fluorescein-labeled naked- or AV-*Bc-DCL1/2*-dsRNA were added to *B. cinerea* spores and fluorescent signals were detected in *B. cinerea* cells after culturing on PDA medium for 10 h. The signals were still visible after adding Triton X-100 to rupture any RNA-containing AVs outside of fungal cells. MNase treatment was performed 30 min before image acquisition. Fluorescence signals of *Bc-DCL1/2-*dsRNA remained visible in *B. cinerea* cells after treated with MNase treatment before observation. Scale bars, 20 μm.

The ability of the AVs to prevent nuclease degradation was then validated under different enzymatic hydrolysis conditions. Naked and AV-loaded *Bc-DCL1/2-dsRNA* were both treated with Micrococcal Nuclease (MNase). As seen in Figure 1B, the naked-*Bc-DCL1/2*-dsRNA exhibited complete degradation after MNase treatment as compared to the AV-protected *Bc-DCL1/2-dsRNA* released from the AV-*Bc-DCL1/2*-dsRNA after the MNase treatment using 1% Triton X-100 that had no obvious degradation. Thus, the AVs provide protection for dsRNA against nuclease degradation.

Finally, we assessed the ability of the AVs as an efficient vehicle for dsRNA delivery to *B. cinerea* fungal cells. Previously, we discovered that naked dsRNA is effectively taken up by *B. cinerea*^7^. Here, we compared fungal uptake of naked and AV-encapsulated Fluorescein-labeled dsRNA using confocal laser scanning microscopy (CLSM). To do this, we placed PDA media directly onto microscope slides, and inoculated with 4 *μ*L of 1×10^5^ spores/mL *Botrytis cinerea*. 4 *μ*L of fluorescein-labeled dsRNA at a concentration of 80 ng/*μ*L was then applied to the slides. Fluorescent dsRNA was detected inside the fungal cells after application of either naked- or AV-*Bc-DCL1* /*2*-dsRNA to *B. cinerea spores* cultured on PDA plates (Figure 1C). The CLSM analysis was carried out after MNase treatment to eliminate any fluorescent signals coming from dsRNA or AV-dsRNA not inside the fungal hyphae. Under these conditions, a strong fluorescent signal was found on and within the hyphaes after AV-dsRNA application, suggesting that the AV-dsRNA were taken up by the fungal cells (Figure 1C).

### External AV-dsRNA application triggers RNAi in *B. cinerea*

After demonstrating that the AVs could be loaded with dsRNA and taken up by fungal cells, we next examined if external AV-dsRNA application triggered RNAi in *B. cinerea*. Naked- and AV-dsRNA were externally applied to a variety of agriculturally relevant plant materials, including tomato and grape fruits, lettuce leaves and rose petals, and a reduction of *B. cinerea* virulence was observed (Figure 2A). These plant materials were treated with 20 μL droplets of the dsRNA treatments, at a concentration of 20 ng/μL. After RNA treatment, 10 μL droplets of fungal suspension (concentration varies and is dependent on different plant material, see methods) were added to the plant material. Two fungal-gene targeting dsRNA sequences were used, both of which have previously been successfully used in SIGS applications against *B. cinerea^7,8^*. One was the above-mentioned BcDCL1/2 sequence, for which we had previously reported that fungal disease was reduced on *Arabidopsis Bc-DCL1/2* RNAi transgenic plants using both host induced gene silencing (HIGS) and SIGS approaches^7^. The other was a sequence of 516 bp containing three fragments of *B. cinerea* genes involved in the vesicle-trafficking pathway: vacuolar protein sorting 51 (*VPS51*, *BC1G_10728)*, dynactin complex large subunit (*DCTN1, BC1G_10508)*, and suppressor of actin (*SAC1*, *BC1G_08464*). These fungal genes were previously described as natural targets in the cross-kingdom RNAi interaction between *Arabidopsis* and *B. cinerea*^37^ and have been proven to be effective for SIGS-mediated inhibition of fungal diseases, such as *B. cinerea*, *Sclerotinia sclerotiorum*, *Aspergillus niger*, *Rhizoctonia solani*, and *Verticillium dahliae ^8^*.Consequently, three dsRNAs were generated by *in vitro* transcription for loading into AVs: two of them specifically targeting *B. cinerea* virulence-related genes (*Bc-DCL1/2* and *Bc-VPS51*+*DCTN1*+*SAC1* (*Bc-VDS*)), while the third one was a non-specific target sequence (*YFP*)used as a negative control. All plant materials treated with naked- or AV-fungal gene targeting-dsRNA (*Bc-DCL1/2* or *-VDS*) had obvious reduced disease symptoms in comparison to the water treatment and *YFP*-dsRNA controls (Figure 2A, 2B). The relative lesion sizes were reduced 75-90%. Further, both naked- and AV-*Bc-VDS* treatments decreased expression of the three targeted fungal virulence genes (Figure S1). Taken together, these results demonstrate how externally applied AV – dsRNA can inhibit pathogen virulence by suppression of fungal target genes. As the SIGS efficacy of *Bc-DCL1/2* and *Bc-VDS* dsRNAs are similar, we used only the *Bc-VDS* dsRNA for the subsequent analysis.

**Figure 2.**
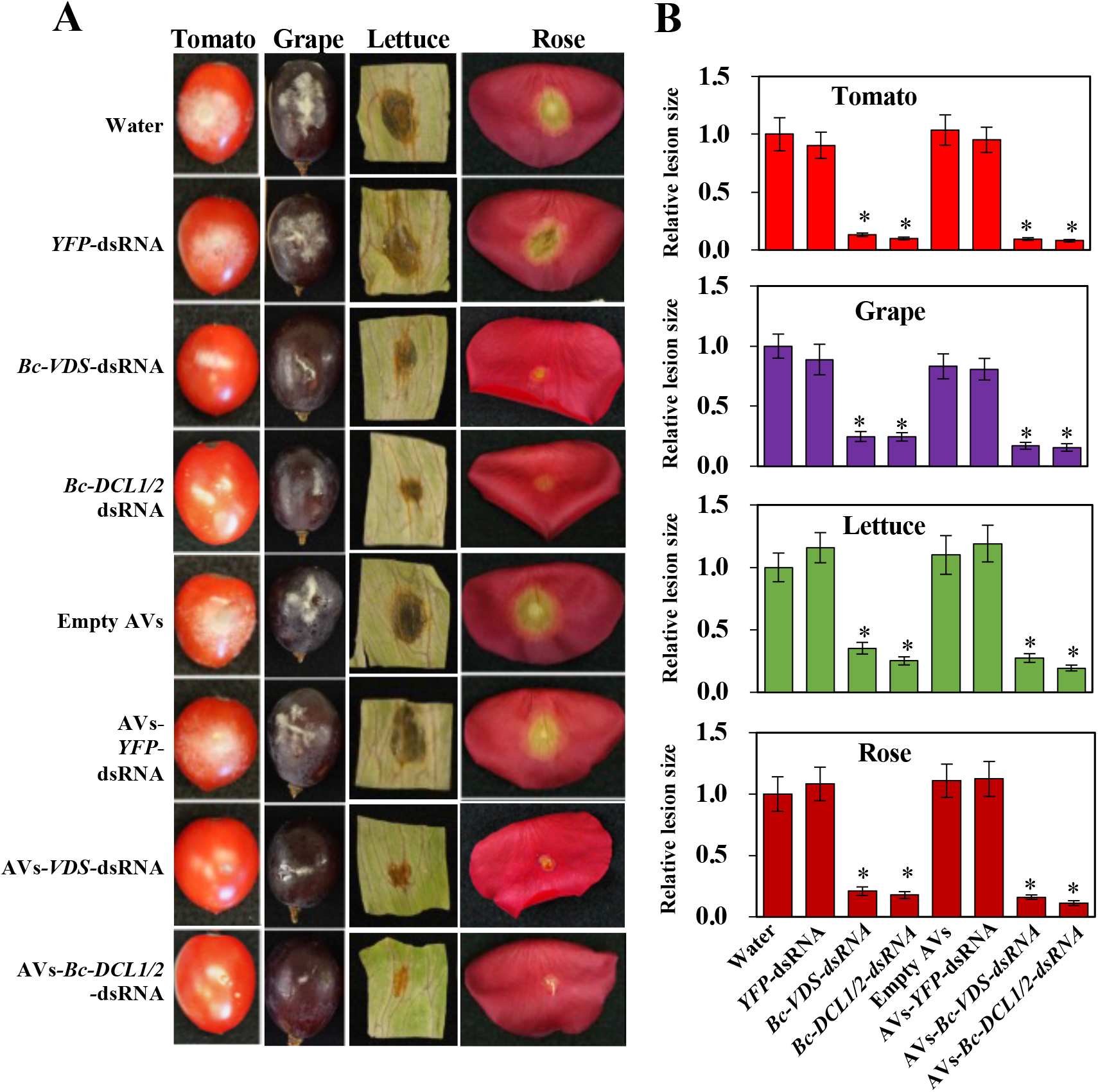
Externally applied naked-dsRNAs or AVs-dsRNA inhibited pathogen virulence. (**A**) External application of naked- and AV-*Bc-VDS*-dsRNA, as well as the application of naked- and AV-*Bc-DCL1/2*-dsRNA, inhibited *B. cinerea* virulence on tomato fruits, grape berries, lettuce leaves and rose petals compared to the water, empty AVs, naked- or AV-*YFP*-dsRNA treatments. These plant materials were purchased from supermarkets and were treated with 20 μL droplets of the dsRNA treatments, at a concentration of 20 ng/μL. After RNA treatment, 10 μL droplets of fungal suspension (concentration dependent on plant material, see methods) were added to the plant material. (**B**) Relative lesion sizes were measured at 5 dpi on tomato and grape berries, and at 3 dpi on lettuce leaves and rose petals, and with the help of ImageJ software. Error bars indicate the SD of 10 samples, and three technical repeats were conducted for relative lesion sizes. Statistical significance (Student’s t-test) compared to water: *, P < 0.05.

### AV-dsRNA extends RNAi-mediated protection against gray mold disease due to increased dsRNA stability and durability

The instability of naked dsRNA currently limits the practical applications of SIGS. Though we demonstrated that AVs can protect dsRNA from nuclease degradation, environmental variables can also influence RNA stability, such as leaf washing caused by rainfall events. Thus, we were interested in evaluating if using the AV-dsRNA would prolong and improve the durability of the RNAi effect on *B. cinerea*.

To assess the influence of washing on the stability and adherence of the AV-dsRNA to plant leaves, we analyzed the intact dsRNA content on the leaf surface using Fluorescein-labeled *Bc-VDS*-dsRNA and Northern blot analysis after water rinsing. The same concentration of Fluorescein-labeled naked- or AV-*Bc-VDS*-dsRNA (20 ng/*μ*L) was applied to the surface of *Arabidopsis* leaves. After 24 h of incubation, the treated leaves were rinsed twice with water by vigorous pipetting. Immediately after, we found that the naked-dsRNA treated leaves showed a drastic decrease in fluorescence compared with AV-dsRNA treated leaves (Figure 3A). These results suggest that most of the naked-dsRNA was washed off, whereas the AV-dsRNA largely remained on the leaves after rinsing (Figure 3A). The effect of the AVs on dsRNA stability over time was also assessed. To do this, we treated 4-week-old *Arabidopsis* plants with Fluorescein-labeled dsRNA or AV-dsRNA, then incubated the plants in dark conditions for 1 or 10 days. We observed a strong fluorescence signal after 10 days on *Arabidopsis* leaves that were treated with Fluorescein-labeled AV-dsRNA, indicating that AVs confer stability to dsRNA (Figure 3B). By contrast, the naked-dsRNA application showed an undetectable fluorescent signal (Figure 3B) and a weak hybridization signal on the Northern blot analysis, compared to AV-*Bc-VDS*-dsRNA treated leaves, which retained *Bc-VDS*-dsRNA (Figure 3C). We further examined whether the AV-dsRNA remained biologically active over time and prolonged protection against *B. cinerea* compared to naked dsRNA. To this end, *Arabidopsis* leaves were inoculated with *B. cinerea* 1-, 3-, and 10-days post RNA treatment (dpt). Both naked and AV-*Bc-VDS*-dsRNA treatments led to a clear reduction in lesion size over the time points assessed (Figure 3D). However, the efficacy of the naked-*VDS*-dsRNA was reduced at a much faster rate than that of the AV-*VDS*-dsRNA, demonstrating that AVs can enhance the longevity of the RNAi effect of the loaded dsRNAs (Figure 3E).

**Figure 3.**
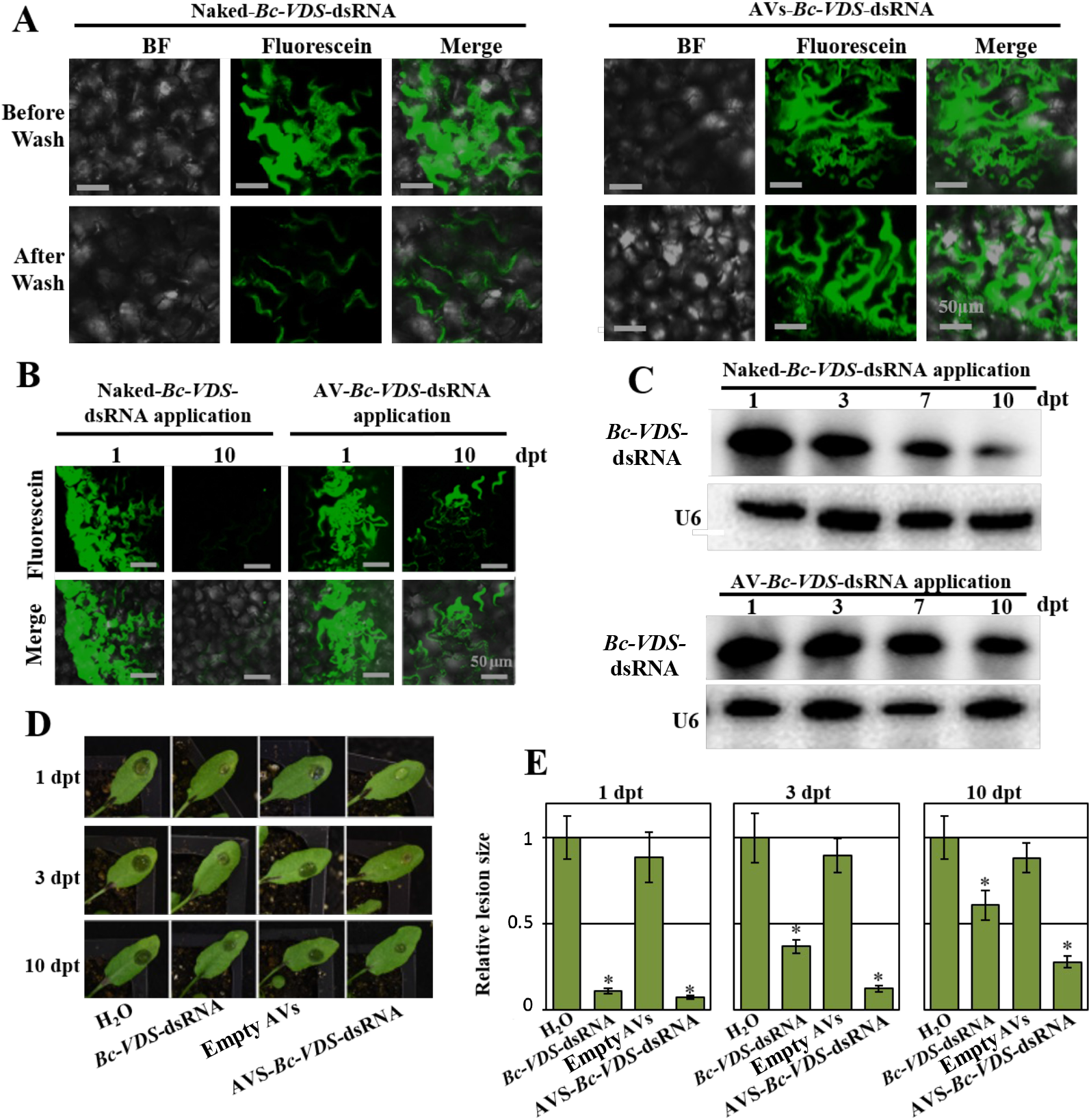
AV-dsRNA has stronger adherence and stability on plant leaves than naked dsRNA. (**A**) CLSM analysis of *Arabidopsis* leaves 1 dpt before and after water rinsing shows the capability of AVs to protect dsRNA molecules from the mechanical action exerted by the water. Scale bars, 50 μm. (**B**) *Arabidopsis* leaves were treated with Fluorescein-labeled naked- or AV-dsRNA for 1 and 10 days. The fluorescent signals on the surface of leaves were observed using CLSM. Scale bars, 50 μm. (**C**) The AV-*Bc-VDS*-dsRNA is highly stable compared with naked *Bc-VDS*-dsRNA on *Arabidopsis* leaves at 10 dpt, as detected by Northern Blot. (**D**) Lesions on *Arabidopsis* leaves inoculated with *B. cinerea* at 1, 3, and 10 dpt. (**E**) Relative lesion sizes were measured 3 dpi with the help of ImageJ software. Error bars indicate the SD. Statistical significance (Student’s t-test) relative to water: *, *P* < 0.05.

To examine if AV-dsRNAs could be similarly effective on economically important crops, we repeated these experiments using tomato fruits, grape berries and grape (*V. vinifera*) leaves. We applied naked- or AV-*Bc-VDS*-dsRNA on the surface of tomato and grape berries and on the surface of grape leaves. The dsRNA was applied via drop inoculation on the fruits, and via spray application on the grape leaves, always at a concentration of 20 ng/*μ*L. Both the naked and AV-*Bc-VDS*-dsRNA applications led to reduced disease symptoms on tomato and grape berries at 1, 5 and 10 dpt, as well as on detached grape leaves at 1, 7, 14 and 21 dpt, compared to the water or empty AV treatments (Figure 4A). As we had observed in the *Arabidopsis* interactions, the AV-*Bc-VDS*-dsRNA applications greatly prolonged and improved the RNAi activity as compared to the naked-dsRNA over time for all plant materials (Figure 4B). While the naked treatment lost the majority of its efficacy at 5-dpt in tomato fruits, 10-dpt in grape berries, and 21-dpt in grape leaves, the AV-dsRNA treatments reduced lesion sizes by at least 75% across all timepoints and plant material tested (Figure 4B). These trends were also reflected in experiments on rose petals after the naked- and AV-*Bc-VDS*-dsRNA treatments (Figure S2). The enhanced reduction in lesion size observed specifically at the longer time points (i.e. 5, 10, 14, and 21 dpt) after AV-*Bc-VDS*-dsRNA application clearly demonstrates how AVs protect loaded dsRNA from degradation to extend the duration of plant protection against *B. cinerea*. Together, these results strongly support the ability of AVs to confer higher RNAi activity over time, effectively enhancing dsRNA stability for SIGS applications.

**Figure 4.**
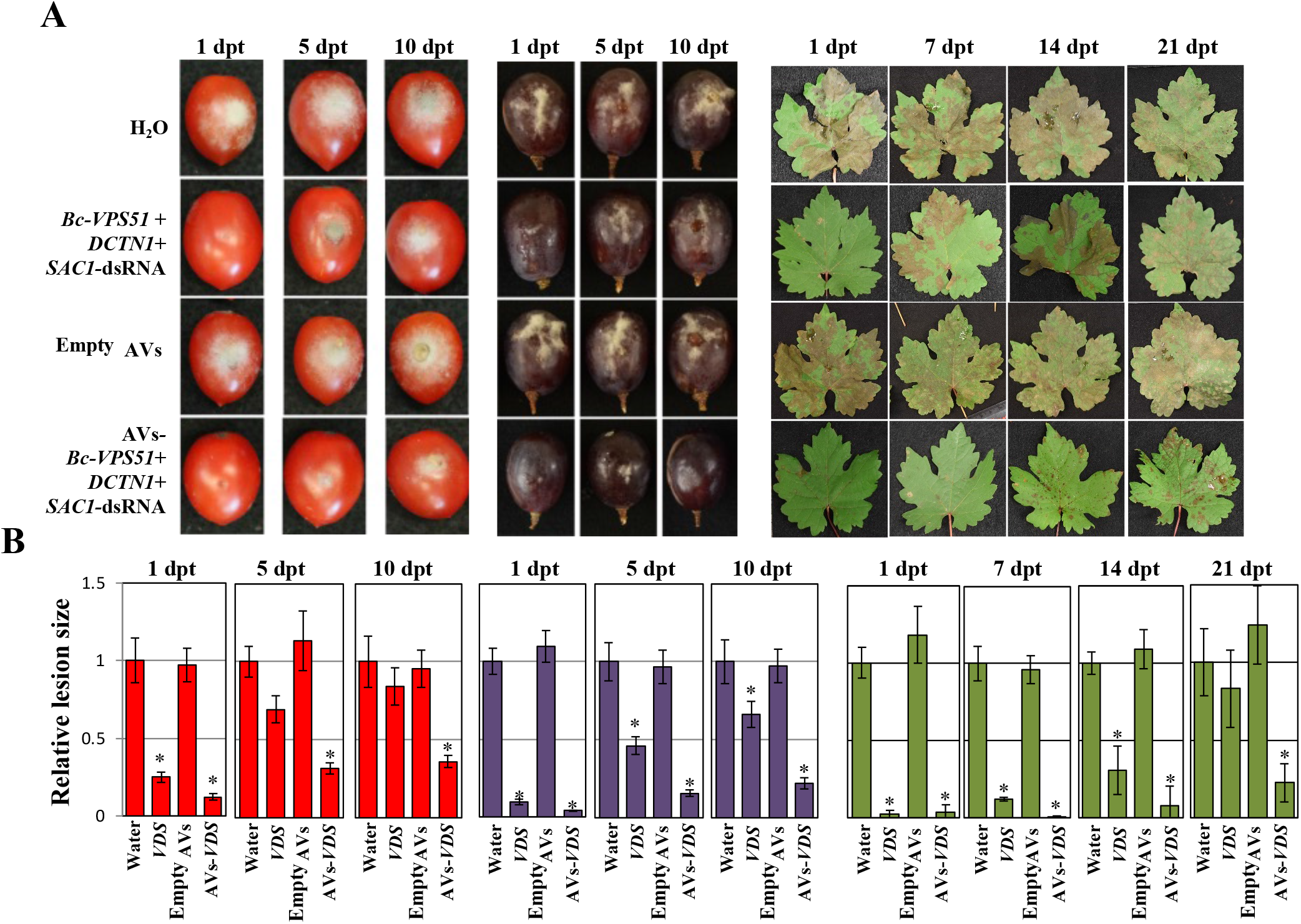
Treatment with AV-dsRNA provides prolonged protection against *B. cinerea* in tomato fruits, grape berries and *V. vinifera* leaves. (**A**) Tomato fruits and grape berries, as well as grape leaves were pre-treated with naked- or AV-*Bc-VDS*-dsRNA, for 1, 5, and 10 days; or 1, 7, 14, and 21 days respectively, then inoculated with *B. cinerea*. Pictures were taken at 5 dpi. (**B**)Relative lesion sizes were measured with the help of ImageJ software. Error bars indicate the SD. Statistical significance (Student’s t-test) relative to water: *, *P* < 0.05.

### Cost-effective AV formulations also provide strong RNAi activity

Our discovery that AVs can lengthen dsRNA mediated plant protection opens the door for its practical use in agricultural applications. Cost is a critical consideration for any crop protection strategy, so we next tested if more cost-effective AV formulations could be used for dsRNA delivery and RNAi activity. First, we removed the PEG, an expensive reagent in the formula, from our original DOTAP+PEG formulation, resulting in DOTAP AVs composed only of DOTAP and cholesterol in a 2:1 ratio. Additionally, we used a cheaper cationic lipid, 1,2-dioleyloxy-3-dimethylaminopropane (DODMA), in a 2:1 ratio with cholesterol to form DODMA AVs. DODMA has previously been utilized in drug delivery formulations, but has a tertiary amine and is an ionizable lipid compared to DOTAP, which could result in changes in RNA loading and activity. The DOTAP AVs were fully loaded with *Bc-VDS* dsRNA at a 1:1 N:P ratio (Figure 5A), requiring the use of 4x fewer lipids than the DOTAP+PEG AVs, or the DODMA AVs, which were completely loaded at a 4:1 N:P ratio (Figure 5B). DOTAP AVs loaded with *Bc-VDS* dsRNA had an average size of 259.5 nm with PDI = 0.368 and ZP = +47.23 mV. These values were similar to those observed for loaded DOTAP+PEG AVs. DODMA AVs loaded with *Bc-VDS* dsRNA though had an average size of 393.6 nm with PDI = 0.385 and ZP = −9.55 mV. Both DOTAP and DODMA formulations could effectively protect *Bc-VDS* dsRNA from nuclease degradation (Figure 5C). The size distribution data for each AV formulation can be found in Table S1. As expected, the z-average sizes of the DOTAP-derived AVs are similar, while the use of DODMA increases the z-average size (Table S1).

**Figure 5.**
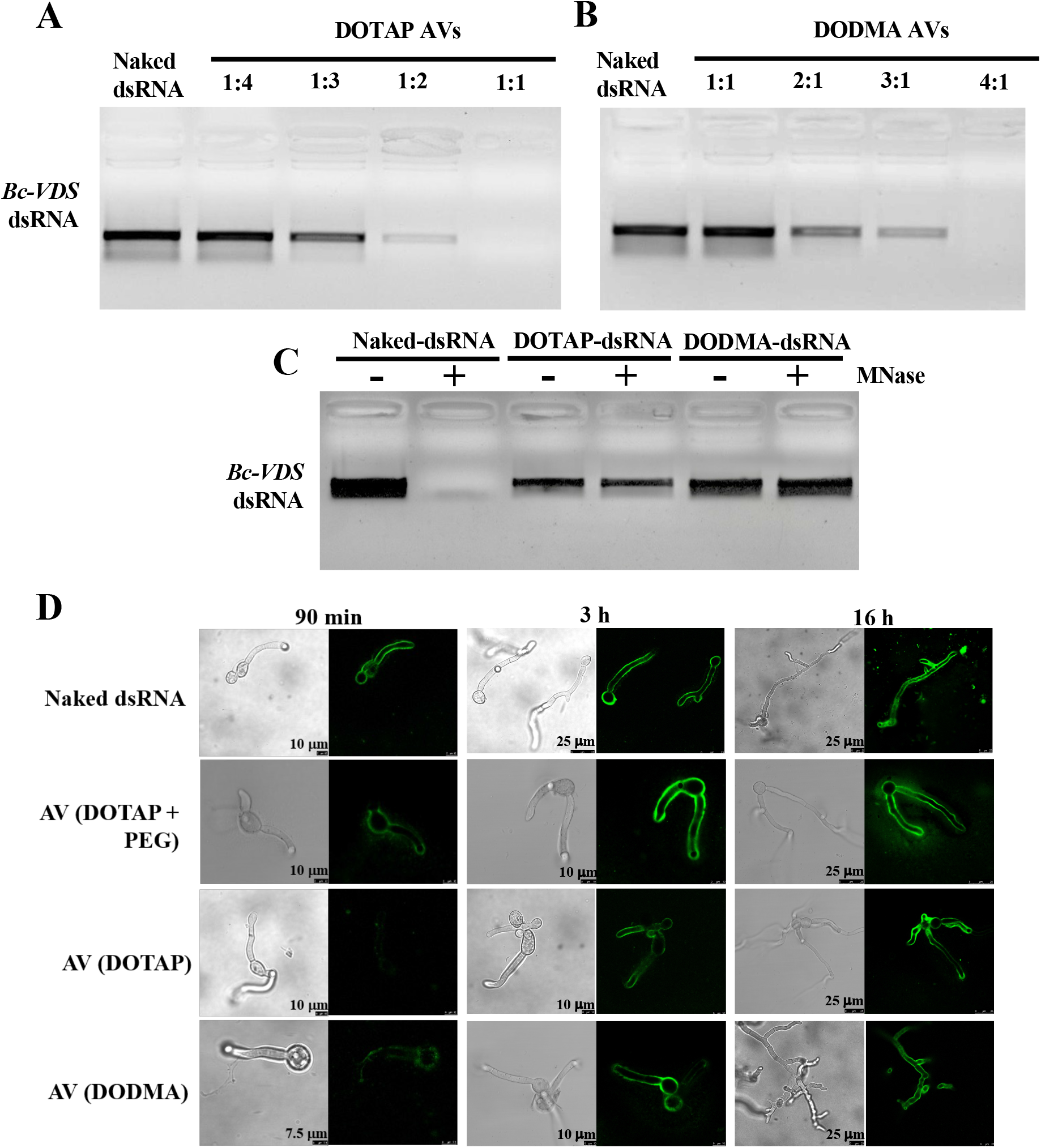
Alternative AV-formulations protect dsRNA from nuclease degradation and are easily taken up by *B. cinerea*. (**A**) DOTAP AV-*Bc-VDS*-dsRNA lipoplexes were formed at a range of indicated charge ratios (N:P) and incubated for 2 h at room temperature before being loaded onto a 2% agarose gel. Complete loading was achieved at an AV:dsRNA mass ratio of 1:1. (**B**) DODMA AV-*Bc-VDS*-dsRNA lipoplexes were formed at a range of indicated charge ratios (N:P) and incubated for 2 h at room temperature before being loaded onto a 2% agarose gel. Complete loading was achieved at an AV:dsRNA mass ratio of 4:1.(**C**) The stability of naked-, DOTAP-, and DODMA-*Bc-VDS*-dsRNA was tested after MNase treatment. *Bc-VDS-* dsRNA was released from AVs using 1% Triton X-100 before gel electrophoresis. (**D**) Analysis of *B. cinerea* uptake of fluorescein-labeled dsRNA encapsulated in three different AV formulations (DOTAP+PEG, DOTAP and DODMA) after 90 mins, 3 and 16 hours of incubation. Fluorescence signals are visible in the *B. cinerea* cells treated with the three AV-*Bc-VDS*-dsRNA using Triton X-100 and MNase treatment before observation.

Next, we examined if the different AV formulations influenced fungal dsRNA uptake or RNAi activity. After application of the different AV formulations, the fungal dsRNA uptake was tracked over 16 hours using CLSM. After 16 hours, all three AV formulations showed a similar amount of fungal RNA uptake, however, the uptake of DOTAP+PEG and DODMA AVs was slightly faster than that of DOTAP AVs without PEG, as evidenced by the stronger signal at the 90 minute and 3-hour timepoints (Figure 5D). To confirm that the lower cost AV formulations have similar RNAi activity on *B. cinerea* over time as our original AV formulation, we performed treatments on tomato fruits purchased from the supermarket. Both the DOTAP and DODMA formulations in complex with *Bc-VDS*-dsRNA trigger a steady RNAi effect on *B. cinerea* over time (Figure 6), significantly reducing lesion sizes at all time points (1, 5 and 10 dpt), in comparison to water. In comparison to the naked *Bc-VDS-*dsRNA, however, only the DOTAP and DODMA formulations significantly reduced lesion sizes at the 10 dpt time point (Figure S3). In addition, fungal biomass quantification indicated that the treatments with *Bc-VDS* dsRNA encapsulated in DOTAP and DODMA AV formulations resulted in a statistically significant reduction of the fungal biomass at all time points. All AV-*VDS*-dsRNA treatments were also able to reduce expression of the targeted *B. cinerea* genes at all time points (Figure S4). Further, no phytotoxicity on *Arabidopsis* plants was observed for any AV formulation, indicating that they can likely be used to protect plants in addition to fruits (Figure S5). Overall, these experiments demonstrate how new AV formulations that are more economical, but equally as effective, can be developed.

**Figure 6.**
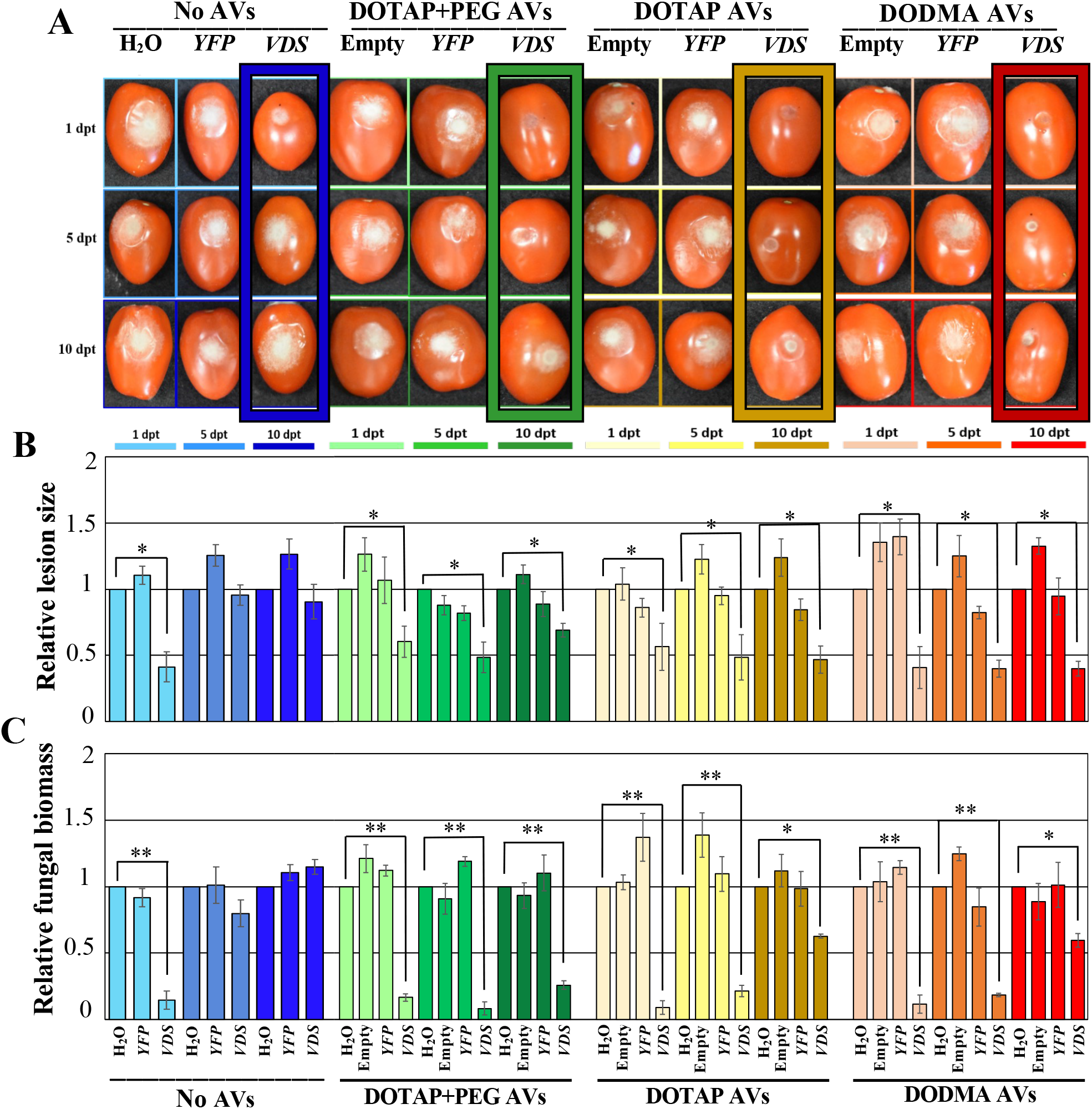
Treatment with all DOTAP+PEG, DOTAP and DODMA AV-dsRNA formulations provide prolonged protection against *B. cinerea* in tomato fruits. (**A**) Tomato fruits were pretreated with naked- or AV(DOTAP+PEG)-*Bc-VDS*-dsRNA, AV(DOTAP)-*Bc-VDS*-dsRNA and AV(DODMA)-*Bc-VDS*-dsRNA, for 1, 5, and 10 days, then inoculated with *B. cinerea*. Pictures were taken at 5 dpi. (**B**) Relative lesion sizes were measured with the help of ImageJ software. Error bars indicate the SD. Statistical significance (Student’s t-test) relative to water: *, *P* < 0.05. (**C**) Relative fungal biomass was quantified by qPCR. Fungal RNA relative to tomato RNA was measured by assaying the fungal actin gene and the tomato tubulin gene by qPCR using RNA extracted from the infected fruits at 5 dpi. Statistical significance (Student’s t-test) relative to water: *, *P* < 0.05; **, *P* < 0.01.

## Discussion

Liposomes have been extensively researched for their applications in clinical contexts^47^, in fact, they have been utilized for drug delivery to human fungal pathogens^51–52^ and are able to transit across the fungal cell wall^53^. Here, we provide the first demonstration that lipid-based nanovesicles can also be used in agricultural contexts, to deliver dsRNA to plant pathogens. The primary advantage that AV-dsRNA offers for SIGS over naked dsRNA is increased dsRNA stability. This is crucial for extending the shelf-life of dsRNA products, since extracellular RNases and other ribonucleases have been identified on fruits and the leaves of important economic crops such as tomato or tobacco^54–55^, and for increasing the length of time needed between RNA applications. In fact, utilizing AV-dsRNAs could extend necessary treatment intervals up to a few weeks, as we demonstrated on both grapes and tomatoes (Figure 4), making SIGS a much more agriculturally feasible crop protection strategy. This is similar to the extended protection provided by inorganic dsRNA complex formulations against viruses on *Nicotiana tabacum* cv. Xanthi leaves and fungal pathogens on tomato plants^27,34^. Another key advantage of utilizing liposome technology for crop protection, especially post-harvest products, is that the success of similar formulations in clinical applications^47^ suggests that the AVs will be safe for human consumption.

With agricultural applications in mind, we tested two more cost effective AV formulations. By removing the PEG from DOTAP-AVs, we can reduce the cost of AV synthesis. PEG is used in liposome preparations in clinical contexts to protect liposomes from immune cell recognition and prolonged circulation time^56^, however, this is not a concern in agricultural applications. Additionally, in our DODMA formulation, we used the lipid DODMA in place of DOTAP, which can further reduce costs. Surprisingly, our DOTAP formulation was able to load dsRNA at a 1:1 N:P ratio, in comparison to a 4:1 N:P ratio observed in other formulations. At this lower loading ratio, the cost of DOTAP AV formulations can be even further reduced. The decreased costs of the DODMA and DOTAP AVs potentially make these formulations more suitable for agricultural use.

In summary, we have provided the first example of utilizing a lipid-based nanoparticle, AVs, for the delivery of dsRNAs in SIGS applications. The AV organic formulations used here confer protection to dsRNA that results in an effective and more durable RNAi effect against the fungal pathogen *B. cinerea* in a wide range of plant products, overcoming the main limitation of SIGS to date. This is one key step forward in the development of RNAi-based fungicides which will help reduce the volume of chemical fungicides sprayed on fields and offer a sustainable option to limit the impact of fungal pathogens on crop production and food security.

## Acknowledgments

We thank Philippe Rolshausen for providing the grape plants. This work was supported by grants from National Institute of Health (R35GM136379), National Science Foundation (IOS 2020731), United State Department of Agriculture (2021-67013-34258) and the CIFAR ‘Fungal Kingdom’ fellowship to H.J.; and two graduate student fellowships, one from National Science Foundation (Research Traineeship grant DBI-1922642) to R.H. in H.J.’s lab, and the other from AMPELOS Grape nurseries organization, Italy to L.C. J.N.S. was also supported by MINECO (PID2019-110459RB-I00), MICINN (PLEC2021-008076) and from the European Union’s Horizon Europe research and innovation programme under the MSCA agreement No 101068728.

## Conflict of interests

The authors declare that they have no competing interests.

## Author contributions

H.J. conceived the idea, designed the experiments and supervised the study. L.Q., R.H., J.N.S., L.C. and A.C. performed the experiments. L.Q., J.N.S, R.H. and H.J. drafted the manuscript. H.J., R.H., J.N.S. and A.C revised the manuscript. All authors read and approved of its content.

## FIGURE LEGENDS

**Supplementary Figure 1. Genes targeted through SIGS-applied dsRNAs are downregulated in *Botrytis cinerea*.** The expression of the *B. cinerea* target genes were down regulated as measured by RT-PCR.

**Supplementary Figure 2. Treatment with AV-dsRNA provides prolonged protection against *B. cinerea* in rose petals**. (**A**)Rose petals were pre-treated with naked- or AV-*Bc-VDS*-dsRNA, for 1, 3, and 7 days, then inoculated with *B. cinerea*. Pictures were taken at 3 dpi. (**B**) The relative lesion sizes were measured with the help of ImageJ software. Error bars indicate the SD. Statistical significance compared to water (Student’s t-test): *, *P* < 0.05.

**Supplementary Figure 3. Direct comparison of AV formulation performance with dsRNA performance.**Tomato fruits were pre-treated with naked- or AV(DOTAP+PEG)-*Bc-VDS*-dsRNA, AV(DOTAP)-*Bc-VDS*-dsRNA and AV(DODMA)-*Bc-VDS*-dsRNA, for 1, 5, and 10 days, then inoculated with *B. cinerea*. Relative lesion sizes were measured with the help of ImageJ software. Error bars indicate the SD of the 3 biological replicates. Statistical significance compared to naked-*Bc-VDS* (Student’s t-test): *, *P* < 0.05.

**Supplementary Figure 4. Treatment with AV-dsRNAs reduces expression of targeted genes in *B. cinerea*.**From tomato treatments shown in Figure 6, relative gene expression of the three targeted genes in *Botrytis, VPS51, DCTN1*, and *SAC1* was quantified by qPCR using RNA extracted from the infected fruits at 5 dpi. Statistical significance compared to water (Student’s t-test): *, *P* < 0.05; **, *P* < 0.01.

**Supplementary Figure 5. AV formulations display no signs of phytotoxicity on *Arabidopsis* plants**.5-week-old *Arabidopsis* plants were treated with a 20 *μ*L suspension on each leaf of either water, DOTAP+PEG-, DOTAP-, or DODMA-AVs. AV treatments were performed at working concentrations or at two times the working concentration. Working concentration is determined by the amount of lipid needed to encapsulate dsRNA at a concentration of 20ng/*μ*L.

**Supplementary Table 1. Size distribution of unloaded and dsRNA-loaded AV formulations**.Size distributions and zeta potential were determined for each of the AV-dsRNA formulations using a Zetasizer Advance.

**Supplementary Table 2. Primers used in this study**.All primers used for constructing DNA templates for the different dsRNAs and for qPCR are listed in the table.

## METHODS

### Plant Materials

Lettuce (iceberg lettuce, *Lactuca sativa*), rose petals (*Rosa hybrida L*.), tomato fruits (*Solanum lycopersicum* cv. Roma), and grape berries (Vitis labrusca cv. Concord) were purchased from a local supermarket. Host plants, including *Arabidopsis thaliana*, tomato (money maker), and grape plants were grown in the greenhouse in a 16/8 photoperiod regime at 24±1°C before use in SIGS experiments.

### *Botrytis cinerea* Culture and Infection Conditions

*B. cinerea* strain B05.10 was cultured on Malt Extract Agar (MEA) medium (malt extract 20 g, bacto protease peptone 10 g, agar 15 g per liter). Fungal mycelia used for genomic DNA and total RNA extraction were harvested from cultures grown on MEA medium covered by a sterile cellophane membrane. For *B. cinerea* infection, the *B. cinerea* spores were diluted in 1% Sabouraud Maltose Broth infection buffer to a final concentration of 10^3^ spores ml^-1^ on tomato leaves and 10^5^ spores ml^-1^ for drop inoculation on the other plant materials^57^, 10 μl of spore suspension was used for drop inoculation of all plant materials used, except tomato fruits, in which 20 μl was used. Infected leaf tissues were cultured in a light incubator at 25 °C for 72 h and fruits for 120 h with constant and high humidity. Fungal biomass quantification was performed following the methods described by Gachon and Saindrenan^58^. The p-values were calculated using Student’s t-test for the comparison of two samples and using one-way ANOVA for the comparison of multiple samples.

### Synthesis and Characterization of Artificial Vesicles

PEGylated artificial vesicles were prepared following previously established protocols^49^. In brief, PEGylated artificial vesicles were prepared by mixing 260 μl of 5% dextrose-RNase free dH2O with the lipid mix and re-hydrating overnight on a rocker at 4°C. The re-hydrated lipid mix was then diluted 4-fold and extruded 11 times using a Mini-Extruder with a 0.4 μm membrane (https://avantilipids.com/divisions/equipment-products/mini-extruder-extrusion-technique).

PEGylated artificial vesicles-dsRNA (20 ng μl^-1^) were prepared in the same manner by adding the appropriate amount of dsRNA to the 5% dextrose-RNase free dH_2_O before combining with the lipid mix. The DOTAP + PEG formulation consisted of 1,2-dioleoyl-3-trimethylammonium propane (DOTAP), cholesterol and 1,2-distearoyl-sn-glycero-3-phosphoethanolamine-N-[methoxy(polyethyleneglycol)-2000] (DSPE-PEG2000) mixed at a 2:1:0.1 ratio, respectively. The DOTAP formulation consisted of 1,2-dioleoyl-3-trimethylammonium-propane (DOTAP), and cholesterol mixed at a 2:1:0.1 ratio, respectively. The DODMA formulation consisted of 1,2-dioleyloxy-3-dimethylaminopropane(DODMA), and cholesterol mixed at a 2:1:0.1 ratio, respectively. All lipids were sourced from Avanti Polar Lipids. The average particle size and zeta potential of the artificial vesicles was determined using dynamic light scattering. All measurements were conducted at 25°C using a Zetasizer Advance instrument (Malvern Instruments Ltd, Malvern, Worcestershire, UK) after 10-fold dilution in filtered Milli-Q water. Data reported is the average of three independent measurements.

### *In Vitro* Synthesis of dsRNA

*In vitro* synthesis of dsRNA was based on established protocols^7^. Following the MEGAscript^®^ RNAi Kit instructions (Life Technologies, Carlsbad, CA), the T7 promoter sequence was introduced into both 5’ and 3’ ends of the RNAi fragments by PCR, respectively. After purification, the DNA fragments containing T7 promoters at both ends were used for *in vitro* transcription. Primers for each DNA fragment can be found in Table S2.

### *In Vitro* Naked- and AV-dsRNA Fluorescence Labeling for Confocal Microscopy

*In vitro* synthesis of dsRNA was based on established protocols^7^. *Bc-DCL1/2*-dsRNA was labeled using the Fluorescein RNA Labeling Mix Kit following the manufacturer’s instructions (MilliporeSigma, St. Louis, MO). For confocal microscopy examination of fluorescent dsRNA trafficking into *B. cinerea* cells, 20 μl of 20 ng μl^-1^ fluorescent RNAs, either naked or loaded into AVs were applied onto 5 μl of 10^5^ spores ml^-1^. Germinating spores were grown on PDA medium and placed on microscope slides. The mycelium was treated with Triton-X, when indicated, and 75 U Micrococcal Nuclease enzyme (Thermo Scientific, Waltham, MA) at 37°C for 30 minutes. The fluorescent signal was analyzed using a Leica SP5 confocal microscope.

### External Application of RNAs on the Surface of Plant Materials

All RNAs were adjusted to a final concentration of 20 ng μl^-1^ with RNase-free water before use. 20 μl of RNA (20 ng μl^-1^) were used for drop treatment onto the surface of plant materials, or, approximately 1 mL was sprayed onto grape leaves before inoculation with *B. cinerea*.

### Stability of dsRNAs Bound to AVs

The potential environmental degradation of dsRNA was investigated by exposure of naked-*Bc VPS51*+*DCTN*+*SAC1*-dsRNA (200 ng) and AV-*Bc-VDS*-dsRNA (200 ng/2.5 μg) to Micrococcal Nuclease enzyme (MNase) (Thermo Fisher) treatment in four replicate experiments. Samples were treated with 0.2 U μL^-1^ MNase for 10 min at 37 °C, and dsRNAs were released using 1% Triton X-100. All samples were visualized on a 2% agarose gel. The persistence of sprayed naked-*BcVDS*-dsRNAs and AV-*Bc-VDS*-dsRNAs (4:1) on leaves was assessed in two replicate experiments by extracting total RNA from leaves followed by a northern blot assay with probes specific to the *Bc-VDS*-dsRNA. 4-week-old *Arabidopsis* plants were treated at day 0 with either a 20 μl drop of *Bc-VPS51*+*DCTN1*+*SAC1*-dsRNAs (20 ng μl^-1^) or AV-*Bc-VDS*-dsRNAs (400:100 ng μl^-1^) and maintained under greenhouse conditions. Single leaf samples were collected at 1, 3, 7, and 10 dpt. Total RNA was extracted using TRIzol and subjected to northern blot analysis.

### Total RNA extraction and qRT-PCR

Plant material infected by *B. cinerea* was collected and frozen at −80°C. Then, total RNA was extracted using TRIzol^™^ Reagent (Thermo Fisher Scientific, Waltham, MA, USA) according to the manufacturer’s instructions. Following extraction, RNA was treated with DNaseI according to the manufacturer’s instructions (Thermo Fisher Scientific, Waltham, MA, USA). We confirmed RNA integrity using 1.2% agarose gels and quantified the amount using a Nanodrop 2000c Spectrophotometer (Thermo Fisher Scientific, Waltham, MA, USA). We confirmed the absence of DNA via the lack of conventional PCR amplification of tomato and *Botrytis tubulin* and *actin* genes, respectively. cDNA was synthesized using the Superscript^™^ III First-Strand Synthesis System (Thermo Fisher Scientific, Waltham, MA, USA) following the manufacturer’s instructions. We used SYBR Green mix (Bio-Rad Laboratories, Hercules, CA, USA) to perform RT-qPCR reactions in a CFX384 (Bio-Rad laboratories, Hercules, CA, USA) with the following thermal profile: 95°C for 15 min, followed by 40 cycles of 94°C for 30 s, 50°C for 30 s, and 72°C for 30 s, and testing single amplification by a specific peak in the dissociation melting curve (0.5°C increments every 10 s from 65°C to 95°C). Fungal biomass was determined based on the relative quantity of *B. cinerea* actin transcript normalized to the *S. lycopersicum* tubulin transcript. Expression of targeted genes was also measured via qRT-PCR, using the same protocol as described above. Expression levels in both gene expression and relative biomass assays were determined via the 2^-ΔΔCt^ method^59^.

## Notes

### Competing Interest Statement

The authors have declared no competing interest.

